# Invasion of forested areas in Gabon (Central Africa) by the Asian tiger mosquito and the potential consequences from the One Health perspective

**DOI:** 10.1101/2022.08.23.504969

**Authors:** Judicaël Obame-Nkoghe, David Roiz, Marc-Flaubert Ngangue, Carlo Costantini, Nil Rahola, Davy Jiolle, David Lehmann, Loïc Makaga, Diego Ayala, Pierre Kengne, Christophe Paupy

**Author notes:** Corresponding authors: Judicaël Obame-Nkoghe, Christophe Paupy.

## Abstract

Since its first record in urban areas of Central-Africa in 2000s, the invasive mosquito, *Aedes albopictus*, has continued to spread across the region, including in remote rural areas, and promoted outbreaks of *Aedes*-borne diseases, such as dengue, chikungunya and Zika. From the One-Health perspective, such invasion might enhance *Ae. albopictus* interactions with wild animals in forest ecosystems and favor the spillover of zoonotic arboviruses to humans. From 2014 to 2018, we monitored the steady spread of this mosquito species in the wildlife reserve of La Lopé National Park (Gabon), and evaluated the magnitude of its colonization of the rainforest ecosystem using ovitraps, larval surveys, BG-Sentinel traps, and human landing catches following an anthropization gradient. We detected *Ae. albopictus* in forest galleries up to 15km away from La Lopé village. However, *Ae. albopictus* was significantly more abundant at anthropogenic sites than in less anthropized areas. The number of eggs laid by *Ae. albopictus* decreased progressively with the distance from the forest fringe up to 200m inside the forest, showing that its occurrence in forest ecosystems is restricted to anthropized-sylvatic interfaces with dense forest. This suggests that *Ae. albopictus* may act as bridge vector of zoonotic pathogens between wild and anthropogenic compartments.

## Introduction

The Asian tiger mosquito *Aedes* (*Stegomyia*) *albopictus* (Skuse, 1894) is one of the most important vectors of arboviruses, such as dengue (DENV), chikungunya (CHIKV) and Zika (ZIKV), which cause diseases of important public health concern, particularly in tropical regions (Paupy et al., 2009). *Aedes albopictus* is an invasive vector species native from Asia that has spread worldwide during recent decades (Battaglia et al., 2016; Benedict et al., 2007; Knudsen, 1995), being estimated that 6.3 billion people currently live in areas suitable for this species increasing the potential arboviral transmission risk (Ryan et al., 2019). Moreover, its progressive expansion around the world (facilitated by global trade) highlights its success as an invasive species, thanks to its capacity to develop desiccation-resistance eggs, to use human-made containers, to develop diapause in temperate areas (Juliano and Lounibos, 2005), and its capacity to adapt to urban, periurban or forested environments where it can persist durably (Fontenille and Powell, 2020).

Where it occurs in Central Africa, *Ae. albopictus* is well established in several urban areas, and its expansion to remote areas (e.g. rural villages) is increasingly reported (Paupy et al., 2012). It might also durably invade forest ecosystems, as suggested by its repeated detection in some forested habitats in continental Africa (Savage et al., 1992). In Central Africa, *Ae. albopictus* was first reported in the early 2000s (Fontenille and Toto, 2001), and has gradually spread across almost the entire region (Bobanga et al., 2018; Ngoagouni et al., 2015; Reis et al., 2017). Now, *Ae. albopictus* is suspected to be the major *Stegomyia* species in Central African urban and rural areas, and a first-line vector for arboviruses (Fritz et al., 2019). For instance, after its introduction in Gabon in 2000s (Coffinet et al., 2007), it rapidly spread across the country, favoring major CHIKV, DENV and ZIKV outbreaks in urban (Caron et al., 2012; Leroy et al., 2009; Pagès et al., 2009; Paupy et al., 2010; Peyrefitte et al., 2008) and in remote rural settings (Paupy et al., 2012). Moreover, because of its capacity to invade new areas, its vector competence for many viruses (Pereira-dos-Santos et al., 2020), its catholic feedings habits that include various animal blood sources, and its capacity to alter the epidemiological equilibrium of arbovirus transmission through the selection of strains associated to enhanced virulence (Vazeille et al., 2007), *Ae. albopictus* might colonize forest ecosystems or their fringes and interact with sylvatic cycles of pathogens. More worryingly, particularly for human health, *Ae. albopictus* presence in forests or at forest margins could favor contacts with potential reservoirs of enzootic arboviruses and thereby promote viral spillover to humans (Pereira-dos-Santos et al., 2020).

During its recent intercontinental range expansion, *Ae. albopictus* has mainly invaded anthropogenic habitats where it can exploit man-made containers and humans as blood source to become an efficient vector for epidemic arboviruses, such as DENV and CHIKV (Rezza, 2014; Staples and Fischer, 2014; Zeller et al., 2016). However, it is not clear whether the species has conserved its ancestral capacity to colonize sylvatic environments, particularly in newly colonized areas in tropical America and Africa (Fontenille and Powell, 2020; Paupy et al., 2009). In Asia, it is currently thought that *Ae. albopictus* is more adapted to forest margins (ecotones), the transition from forest to degraded or secondary forests, and open grasslands or scrub rather than to deep primary forests (Fontenille and Powell, 2020). Recent studies confirmed that *Ae. albopictus* has retained its ancestral capacity to colonize forest environments by laying eggs in natural breeding sites and by feeding on non-domestic vertebrates (Pereira-dos-Santos et al., 2020). In Brazil, *Ae. albopictus* was observed up to 500 m from the forest margins and may be established in a degraded forest of the Manaus region (Pereira dos Santos et al., 2018). Despite its detection in forests bordering human villages in (Paupy et al., 2012; Savage et al., 1992), the penetration and invasion dynamic of *Ae. albopictus* in forest ecosystems and more importantly, its persistence as sylvatic populations (i.e. in the absence of humans) have never been studied in Africa.

In Central Africa, natural parks are among the potential invasion territories for *Ae. albopictus*. These parks attract many visitors and are potential hotspots of arbovirus transmission from wildlife reservoirs to humans by zoo-anthropophilic mosquitoes, such as *Ae. albopictus*. In Gabon, La Lopé National Park (LNP) includes large forest-savanna mosaic areas and primary forest blocks, and might be a hotspot of arbovirus circulation (Table 1). Therefore, it is a suitable field of investigation to test whether *Ae. albopictus* can enter and settle permanently in forest ecosystems, through interactions with the wild fauna, and whether it represents a risk of zoonotic arbovirus transfer to humans. To test this hypothesis, in this study carried out at La Lopé village and the northern part of the adjoining the LNP, we monitored *Ae. albopictus* invasion of the wild compartment. We aimed to assess its degree of penetration into the forest ecosystem, from the borders forming anthropized-sylvatic and forest-savanna interfaces, towards the deeper parts of the forest compartment.

**Table 1:**
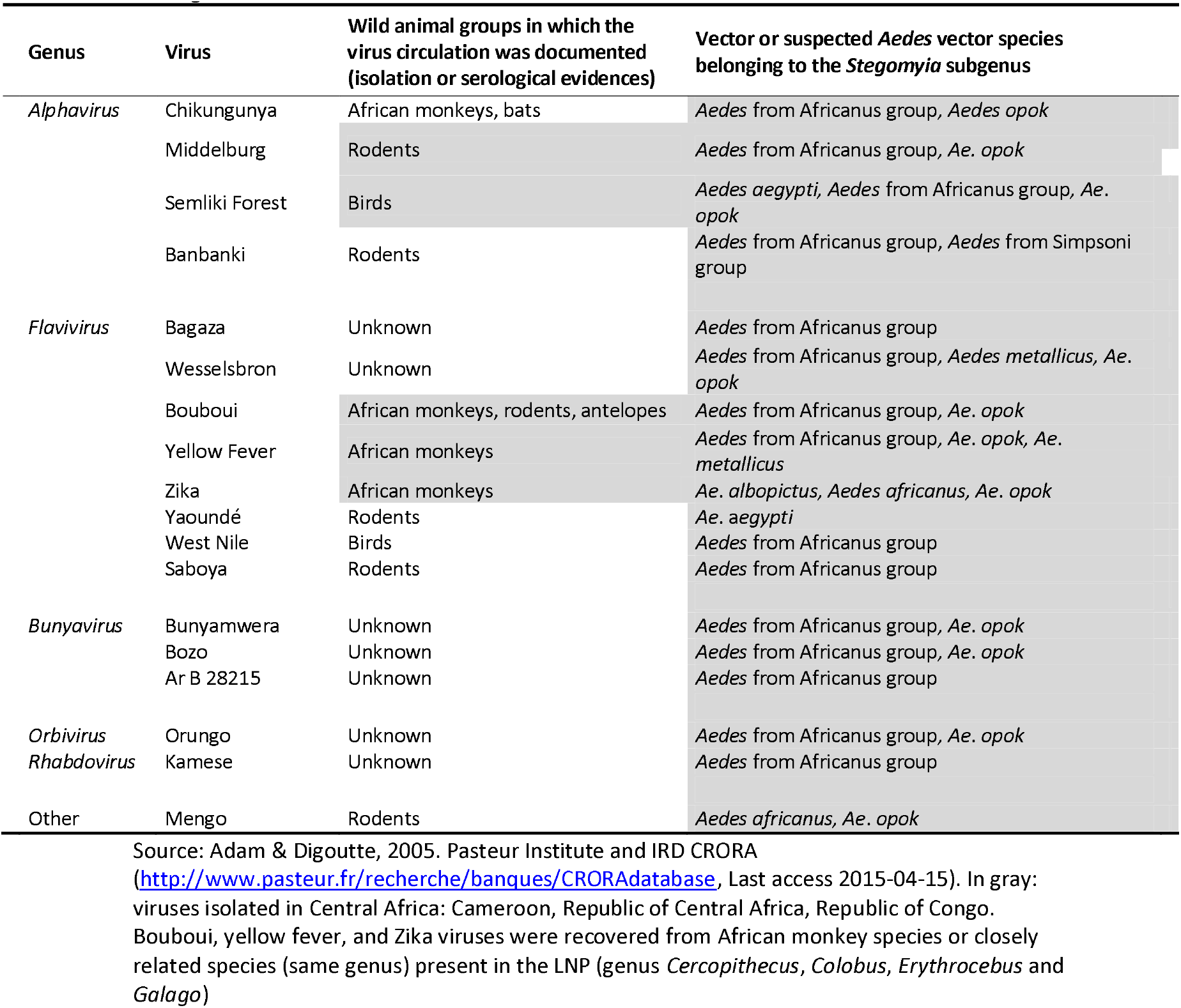
Arboviruses isolated from sylvatic *Stegomyia* species in Central Africa and possibly circulating in the LNP

## Results

### *Aedes albopictus* persistent invasion of uninhabited environments surrounding La Lopé village

From 2014 to 2018 we investigated the range of dispersion of *Ae. albopictus* in La Lopé village and the wild compartment of the LNP using ovitraps and BG-Sentinel traps. For ovitraps, we randomly selected 35 sampling sites, 10 sites in the village and 25 sites southwards in the wild compartment (Figure 1). We collected *Ae. albopictus* eggs at all 10 collection points monitored in the village in 2014 (100%), confirming that the species was well established and widely distributed across the village (i.e. human compartment) (Figure 1). Moreover, adult collections with BG-Sentinel traps in the village (1,340 hours of sampling effort) showed that *Ae. albopictus* was certainly the dominant species compared with other native *Aedes* species from the *Stegomyia* subgenus (Table 2). In addition, the results of human landing collections confirmed that *Ae. albopictus* is the most abundant diurnal mosquito feeding off humans. In the sylvatic compartment, 20 of the 25 ovitrap collection sites surveyed in 2014 (80%) were positive for *Ae. albopictus* (Table 2). This clearly shows that *Ae. albopictus* had widely spread throughout the northern part of the LNP, including sites with few or no human presence. The 20 positive sites were along interconnected forest galleries (n=10), in forest patches (n=4), and at deep forest fringes (n=6) over large expanses in the national park, ranging from 1 to 15 km away from the village (Figure 1). Occasional observations in 2017 and 2018 confirmed the persistence of *Ae. albopictus* over several years in the sylvatic compartment (Figure 1).

**Table 2:** Abundance and percentage of Aedes mosquito species from the Stegomyia subgenus collected in the LNP in 2014 using BG-Sentinel and Human landing catches sampling techniques.

|  | Anthropic compartment |  |  | Sylvatic compartment |  |  | Both compartments |
| --- | --- | --- | --- | --- | --- | --- | --- |
|  | BG-Sentinel (%) | HLC (%) | Sub-total (%) | BG-Sentinel (%) | HLC (%) | Sub-total (%) |  |
| **Invasive species** |  |  |  |  |  |  |  |
| *Ae. albopictus* | 482 (56.1) | 126 (87.5) | 608 (60.6) | 110 (39.1) | 126 (88.1) | 236 (55.7) | 844 (59.1) |
| **Native species** |  |  |  |  |  |  |  |
| *Ae. aegypti* | 80 (9.3) | 18 (12.5) | 98 (9.8) | 20 (7.1) | 15 (10.5) | 35 (8.3) | 133 (9.3) |
| *Ae. Africanus* group | 10 (1.2) | 0 | 10 (1) | 13 (4.6) | 0 | 13 (3.1) | 23 (1.6) |
| *Ae. Apicoargenteus* group | 4 (0.5) | 0 | 4 (0.4) | 7 (2.5) | 0 | 7 (1.6) | 11 (0.8) |
| *Ae. Dendrophilus* group | 10 (1.2) | 0 | 10 (1) | 21 (7.5) | 0 | 21 (4.9) | 31 (2.2) |
| *Aedes fraseri* | 2 (0.2) | 0 | 2 (0.2) | 11 (4) | 2 (1.4) | 13 (3.1) | 15 (1.0) |
| *Ae. (Stegomyia)* spp. | 271 (31.5) | 0 | 271 (27) | 99 (35.2) | 0 | 99 (23.3) | 370 (26.0) |
| **Total** | **859 (100)** | **144 (100)** | **1003 (100)** | **281 (100)** | **143 (100)** | **424 (100)** | **1427 (100)** |

HLC: Human Landing Catch; Sampling efforts: BG-Sentinel (1,340 hours and 1,645 hours in anthropic and sylvatic compartments, respectively) and HLC (6 hours and 12 hours in anthropic and sylvatic compartments, respectively).

**Figure 1:**
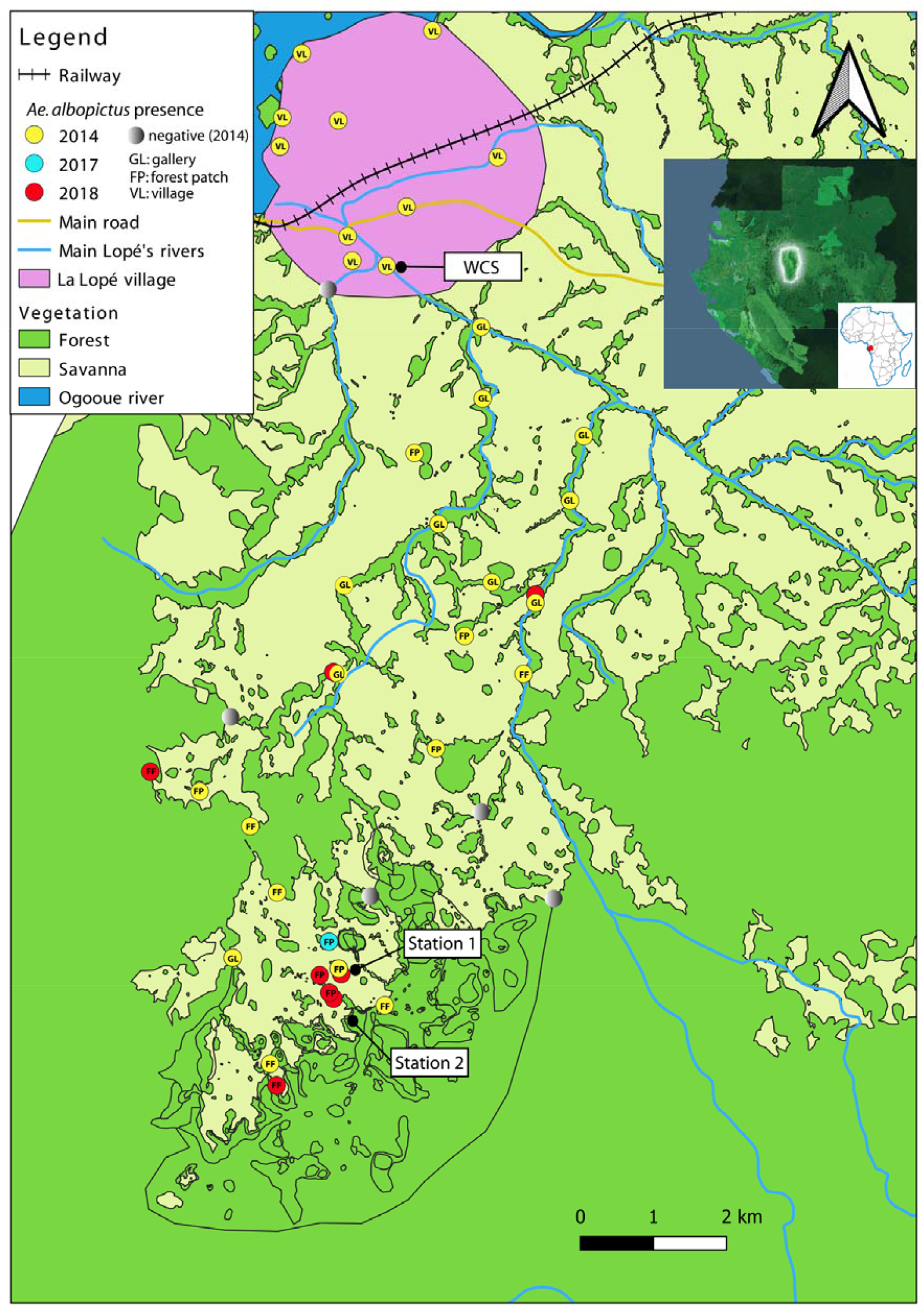
Study area and sampling sites with *Ae. albopictus* occurrences recorded between 2014 and 2018 in the village and in the sylvatic compartments of La Lopé National Park between 2014 and 2018. Sites with *Ae. albopictus* occurrences are marked with yellow, blue, and red circles for 2014, 2017, and 2018, respectively. Black circles indicate the *WCS, Station 1* and *Station 2* sites where ovitrap-based experiments were carried out along anthropization gradients to assess *Ae. albopictus* penetration pattern inside forest ecosystems.

In the sylvatic compartment, *Ae. albopictus* represented 37.2% of all collected *Stegomyia* species using BG-Sentinel traps (110 *Ae. albopictus* among the 296 *Stegomyia* specimens collected during 1,340 hours of sampling; Table 2), and 86.9% of all human-biting mosquitoes from this subgenus (252 *Ae. albopictus* among the 290 *Stegomyia* specimens collected; Table 2). This indicated that *Ae. albopictus* conserved its human feeding habits, even after invading the sylvatic environment where the human presence is limited or absent. It also confirmed the presence of *Ae. albopictus* in forest galleries and at the forest fringes in the northern part of the LNP, raising the question of its capacity to colonize areas deeper in the LNP primary forest ecosystems.

### *Aedes* albopictus relative abundance in relation to distance from the forest ecotone

To determine *Ae. albopictus* colonization level in the forest ecosystem, in 2019, we collected, using plastic ovitraps, mosquito eggs at three ecotone sites with contrasting characteristics (i.e distance from the village, level of anthropization) along forest penetration transects (Figure 1 and Table 3). Out of 4,940 collected eggs over all sites, 2,467 were collected at *WCS* (49.9%), 1,831 at *Station 1* (37.0%), and 642 at *Station 2* (13.1%). After egg hatching and larval development, 2,415 adult mosquitoes emerged and then morphologically identified. The global emergence success from egg to adult stage was 48.9%, but varied significantly according to the collection site (Chi-squared = 165.1, df = 2, *p* < 0.001): 58% for *WCS*, 49% for *Station 1*, and 14 % for *Station 2*. Potential uncontrolled artifacts of conservation or larval growth in plastic trays might explain these differences. Out of the 2,415 emerged adults, 1,431 (59.2%) came from *WCS*, whereas 893 (36.9%) and 91 (3.9%) came from *Station 1* and *Station 2* respectively. A generalized mixed effects model in which the percentage of adult emergence (all species included) was the response variable showed that there was no effect of “Distance from the forest edge” (fixed effect) (*p* = 0.7), regardless of the sampling site (random effect). This suggests that the percentages of adults at the three sites were not skewed, regardless of the sampling site or the penetration depth into the forest (which could reflect a realistic diversity and abundance of mosquitoes that exploited ovitraps deployed at the three sites).

**Table 3:** Eggs collected and adult emergence percentages at the three egg sampling sites

| Site | Total egg number | Number of adults and percentage (%) |  |  | WNAE* |
| --- | --- | --- | --- | --- | --- |
|  |  | *Ae. albopictus* | Other species | Total |  |
| **WCS** | 2467 | 1187 (82.9) | 244 (17.1) | 1431 (100) | 2045 |
| **Station 1** | 1831 | 525 (58.8) | 368 (41.2) | 893 (100) | 1075 |
| **Station 2** | 642 | 9 (9.9) | 82 (90.1) | 91 (100) | 63 |

WNAE: Total Weighted Number of *Ae. albopictus* eggs

**Table 4:** Ovitraps colonized by *Ae. albopictus*

| Site | Habitat | N. of traps deployed | N. of positive traps | % | % C-interval |
| --- | --- | --- | --- | --- | --- |
| WCS | Gallery | 35 | 24 | 68.6 | [50.7-83.1] |
| Station 1 | Patch | 29 | 26 | 89.6 | [72.6-97.8] |
| Station 2 | Forest | 31 | 2 | 6.5 | [0.7-21.4] |

C-interval: Confidence interval of positive trap percentage; %: Percentage of positive traps

Regarding the species relative abundance (determined as species percentage) according to the distance from the forest edge and sampling site (representing three levels of anthropization), we observed that *Ae. albopictus* was the dominant species at *WCS* (high anthropization) at all distances except at 100m and 150m from the forest edge. It was followed by *Aedes* from the Africanus group (*Ae*. gr. Africanus) for which the relative abundance increased with the distance from the forest edge, and the highest relative abundances were at 100m and 150m from the forest edge (Figure 2A). At *Station 1* (low anthropization), *Ae. albopictus* was dominant from 0m to 50m and at 100m from the forest edge, followed by *Ae*. gr. Africanus that displayed the highest relative abundances at 75m and 125m from the forest edge (Figure 2A). Conversely, at *Station 2* (no anthropization), *Ae*. gr. Africanus was the most abundant, regardless of the distance from the edge, but not at 25m (Figure 2A). In all three sites, *Ae. albopictus* and *Ae*. gr. Africanus were the two predominant species. However, their relative abundance varied according to the site and the degree of wilderness. Indeed, the relative abundance of *Ae. albopictus* decreased with increasing wilderness (p = 0.04), whereas the relative abundance of *Ae*. gr. Africanus increased with wilderness (p < 0.001) (Figure 2B).

**Figure 2:**
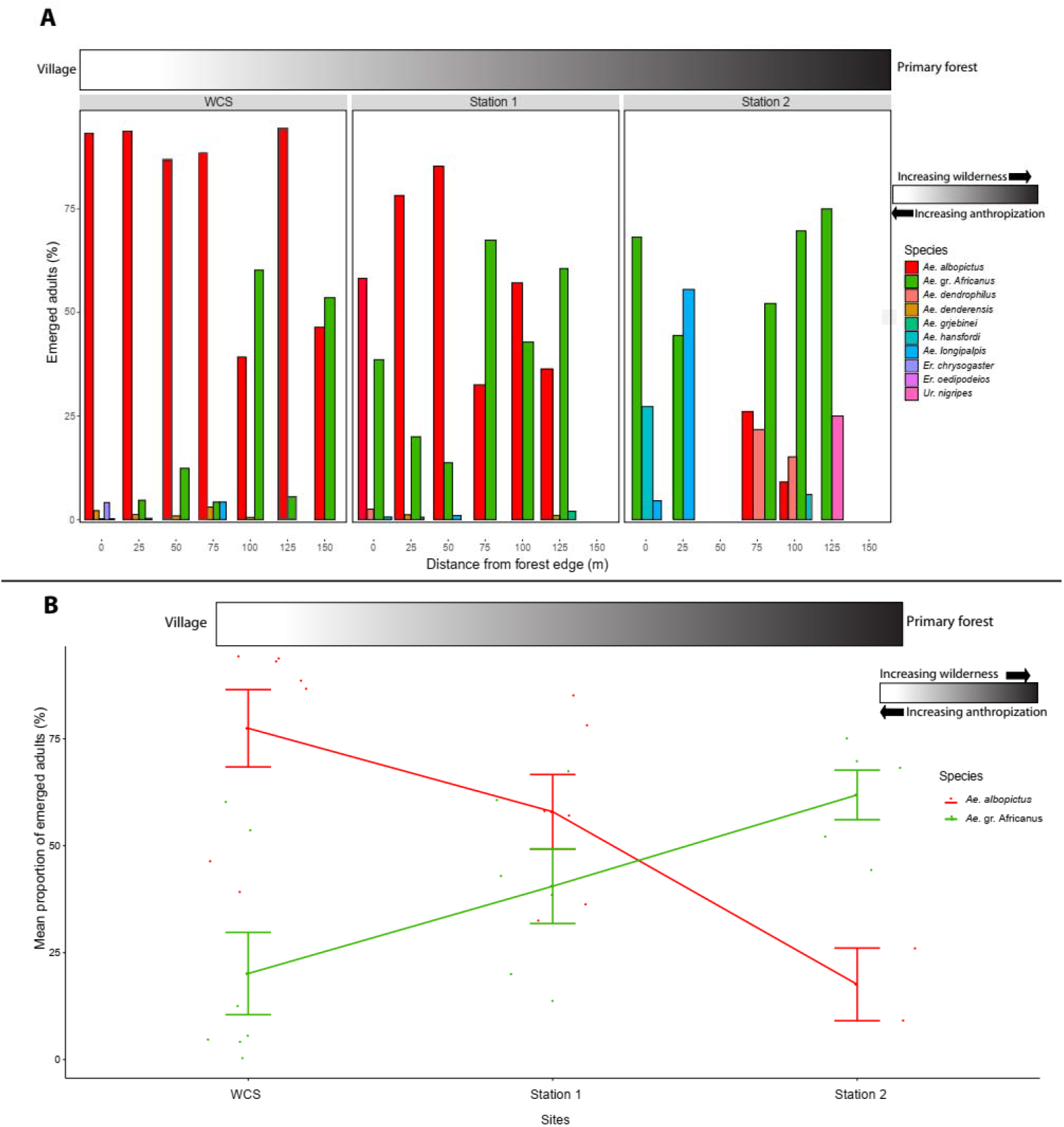
Relative abundance of *Ae. albopictus* and other species recovered from ovitraps distributed according to a gradient of anthropization from village to primary forest in Lopé National Park, Gabon. **A:** Relative abundance of all species. **B**: Mean percentage of emerged adults for the two predominant taxa: *Ae. albopictus* and *Ae*. gr. Africanus in three sites arranged according to degree of anthropization.

When we analyzed adult stages that emerged from collected eggs, we observed a significant difference (Pearson’s Chi-square = 330.6, df = 2, *p* < 0.001) in the percentage of *Ae. albopictus* specimens for *WCS* (82.9%), *Station 1* (58.8%), and *Station 2* (9.9%) (Table 3). Moreover, the percentage of *Ae. albopictus* positive ovitraps differed significantly at the three sites (Pearson’s Chi-square = 46.1, df = 2, *p* < 0.001). The findings indicate that at *WCS* and *Station 1* ovitraps were colonized in similar proportions (Pearson’s Chi-square = 2.9, df = 1, p = 0.08), suggesting that *Ae. albopictus* presence was more associated with sites showing anthropogenic features.

We then used the percentage of adult specimens to calculate the estimated total weighted number of *Ae. albopictus* eggs collected in each site (WNAE, defined as the frequency of *Ae. albopictus* emerged adults multiplied by the total number of eggs collected) (Table 3). Comparison of the WNAE values showed that *Ae. albopictus* was the predominant species at sites with substantial anthropization (*WCS* and *Station 1*: characterized by the presence of human habitations with domestic water containers, tires, metallic and plastic containers from human activities), whereas it was weakly represented at the uninhabited site (*Station 2*). When we compared the mean number of *Ae. albopictus* eggs (i.e. the mean number of *Ae. albopictus* eggs laid in each ovitrap) among sites, we observed that this number significantly decreased (Kruskal-Wallis chi-squared = 23.79, df = 2, *p* < 0.001) from *WCS* (highly anthropized forest edge) to *Station 2* (lowly anthropized forest edge) (Figure 3).

**Figure 3:**
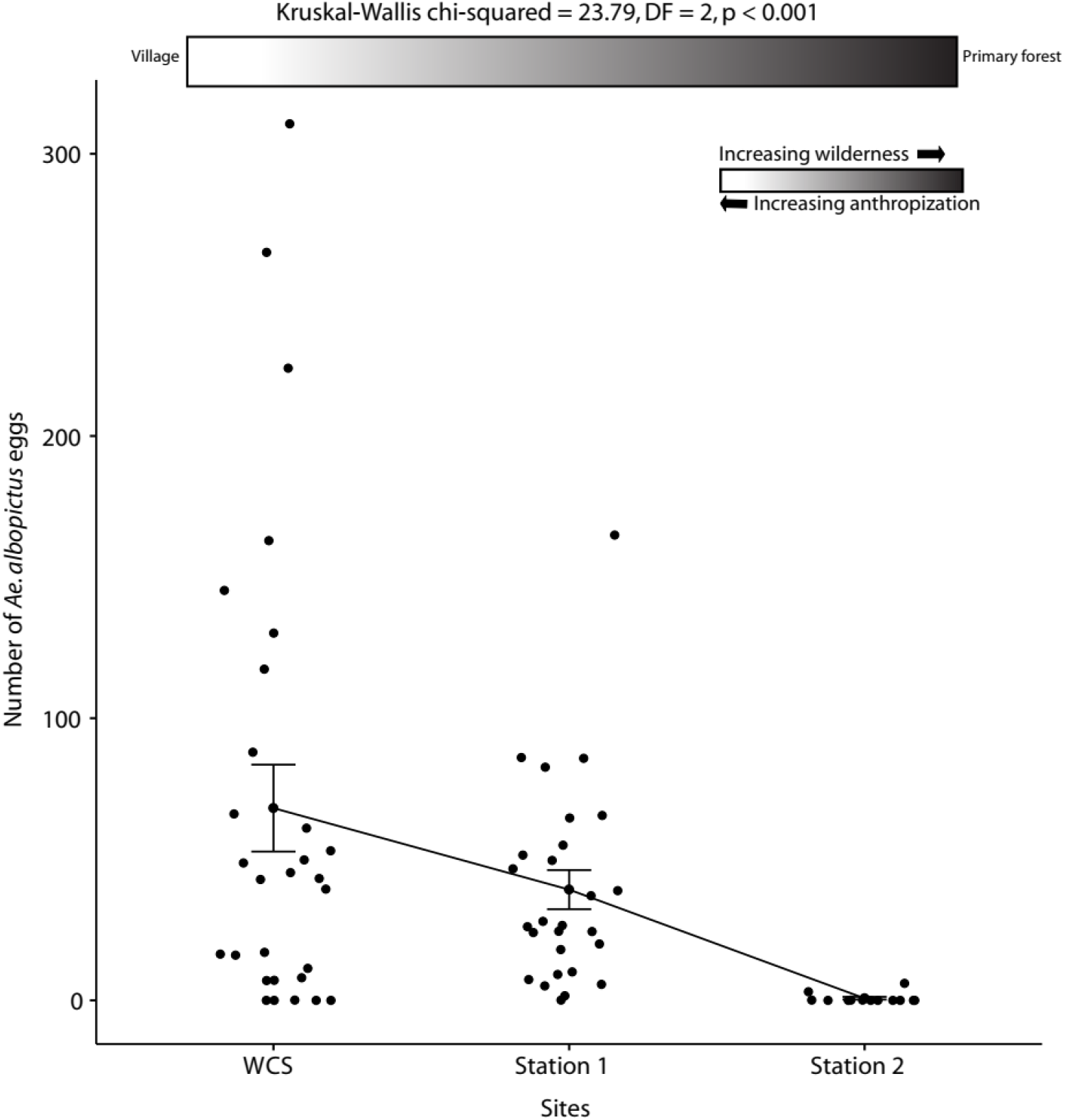
Comparison of the number of *Ae. albopictus* eggs laid in ovitraps following a decreasing gradient of anthropization from *WCS* (many human habitations) to *Station 2* (no human habitation).

### Model of *Ae. albopictus* penetration into the forest ecosystem

We evaluated *Ae. albopictus* colonization potential of the forest ecosystem by analyzing its oviposition activity at increasing distance from the forest fringe. The Shapiro-Wilk normality test (Figure S1) showed that the WNAE values did not follow a normal distribution (W = 0.75, *p* <0.001). We tested four different generalized linear (mixed) models with residuals following either the Poisson or the negative binomial distribution, and the “Habitat” variable included as a random effect, or otherwise without random effects. After model testing, the Poisson models were discarded because they showed data over-dispersion, thus we focused on the negative binomial negative models. The best model was selected based on Likelihood Ratio tests (LR). The LR test indicated that random effects (*p* = 0.0002) and the “Distance” to the forest edge (z-value = -3.7, df = 52, *p* < 0.001) must be included in the minimal adequate model. The model parameters indicated that the number of *Ae. albopictus* eggs detected in the forest decreased progressively with the distance to the forest edge at both *Station 1* and *WCS* (Figure 4A). At *Station 1*, the mean number of eggs per trap decreased significantly from >50 at the forest edge to <10 beyond 100 meters from the forest edge (Figure 4A), with density peaks spatially concentrated along the forest edge (Figure 4B). Similarly, at *WCS*, the number of *Ae. albopictus* eggs decreased from >200 at the forest edge to <25 after 225 meters from the forest edge (Figure 4A), with density peaks spatially concentrated along the forest edge (Figure 4B). However, at *Station 2*, with only nine specimens recovered (3 at 50m and 6 at 75m from the forest margin), no clear pattern could be deducted. These data showed that inhabited areas of forest fringes are the main *Ae. albopictus* entry points into wild forest ecosystems.

**Figure 4:**
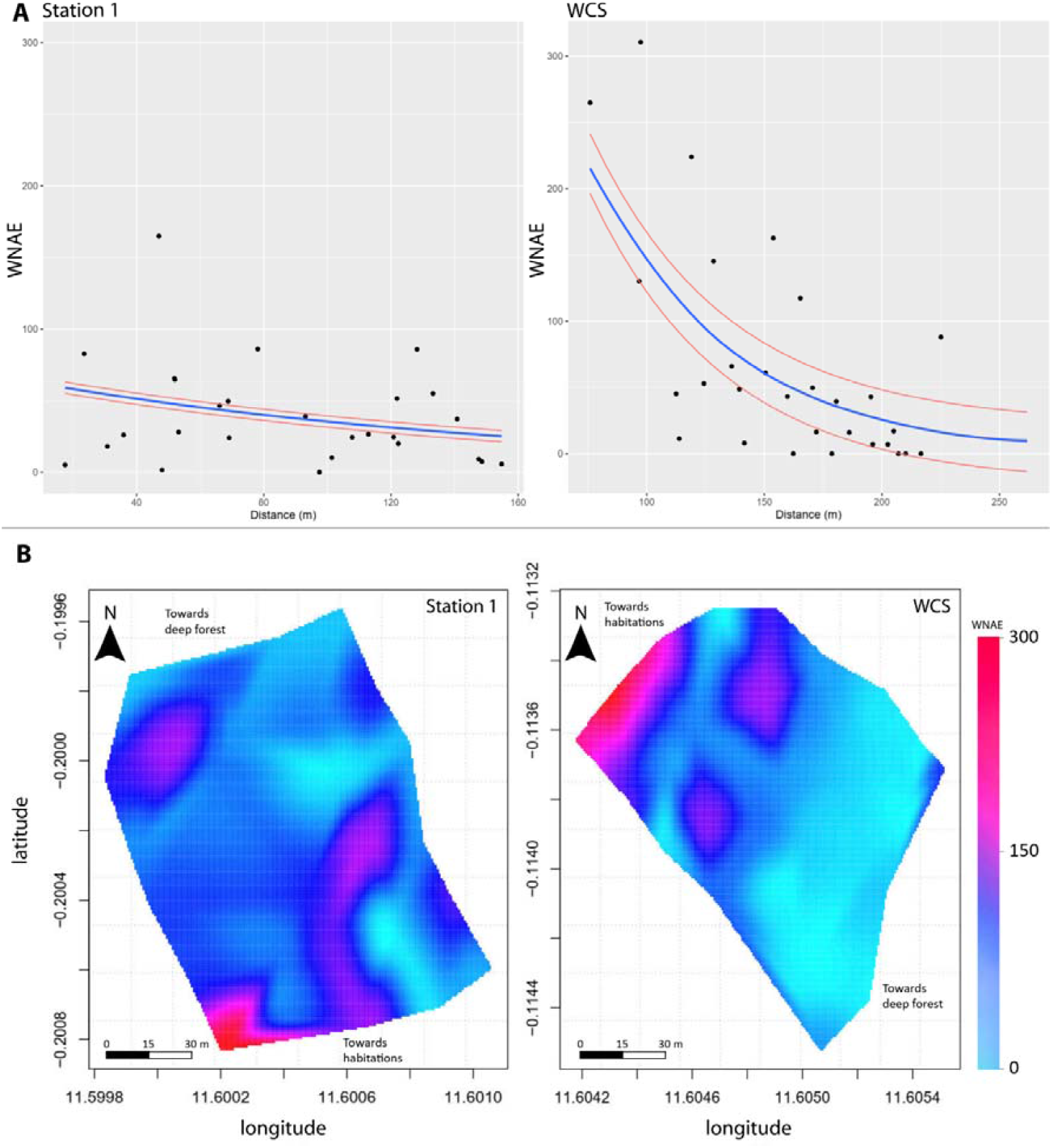
Fitting models of *Ae. albopictus* penetration in the forest ecosystem from the margins of anthropized-sylvatic interfaces. **A**: Scatterplots of the number of *Ae. albopictus* eggs as a function of distance from the forest fringe. Dots indicate observed values and curves show the fitted Negative Binomial regression lines (± 95% confidence limits). The number of eggs in ovitraps progressively decreases with distance from the forest margin. **B:** Two-dimensional heatmap of *Ae. albopictus* egg density at *Station 1* and *WCS* showing density peaks at the forest margins close to human habitations. Latitude and longitude coordinates are expressed according the WGS 84 spatial reference system.

## Discussion

### Occurrence of sylvatic populations of *Aedes albopictus*

To the best of our knowledge, we report here for the first time a situation where *Ae. albopictus* occurs into the wild environment in continental Africa as for native populations in South-East Asia. Indeed, the multiple occurrences of this species at diverse locations throughout the sylvatic compartment of the LNP, up to 15 km from the village, and its persistence over several years at sites distant from human habitations, indicates that these populations might persist in sylvatic populations in areas where the human footprint anthropogenic influence varies from moderate to absent. The human footprint here is mainly limited to ecotourism and associated activities.

In the northern part of the LNP, we recorded *Ae. albopictus* occurrence at many sites located mostly in forest corridors across the savanna interconnecting the village to the primary forest at multiple locations southwards. These forest galleries, rarely exceeding 200 meters in width, may be highly suitable for this species, as previously described in similar environments in its native range in Asia (Fontenille and Powell, 2020) and in South-West islands of the Indian Ocean, such as La Réunion (Delatte et al., 2008). Potential natural larval breeding sites (rock holes and tree holes) and wild animal (as a blood-source for adults) are abundant in these galleries. Thus, it seems likely that *Ae. albopictus* gradually and actively progressed along these forest galleries from the village towards the primary forest. Moreover, its occurrence in fragments of isolated forest patches suggests that this mosquito actively crosses the savanna to colonize new areas, and/or disperse passively. Human activities in the park (e.g. ecotourism, conservation, and research) requiring car shifting from/to the village are an very effective means for passive displacement of *Ae. albopictus* adults even over long distances (Eritja et al., 2017).

At La Lopé village (i.e. the anthropic compartment), the essential resources needed for *Ae. albopictus* life cycle are mainly provided by humans (i.e. blood sources and artificial water containers for larval development). However, as human presence decreases, the need to exploit alternative resources (i.e. wild animals and natural water containers) increases until it becomes the only option at sites without anthropogenic resources. The latter situation was observed at many sites throughout the northern part of the LNP, suggesting that *Ae. albopictus* exploited its opportunistic ecological preferences (Pereira Dos Santos et al., 2020) to recover an ancestral sylvatic ecology (Fontenille and Powell, 2020). Little is known about *Ae. albopictus* forest ecology in Africa, but we observed (unpublished data) that natural breeding sites suitable for this species (mainly water-filled tree and rock holes) are abundant also in the forested environment of the LNP, in dense (e.g. galleries along rivers) and patchy (e.g. in forest-savanna mosaic) areas. Nevertheless, the successful colonization of such natural microhabitats implies that *Ae. albopictus* has overcome several major biotic and abiotic constraints, demonstrating its adaptability. Biotic constraints include larval competition for space and food with the numerous resident mosquito species and other competing aquatic microorganisms. In Africa, tree holes host diversified communities of larval mosquitoes (Dunn, 1927; Pajot, 1983; Service, 1965) that are shaped by interspecific competition, leading to species segregation in space and time to minimize niche overlap (Lounibos, 1981). The colonization of tree-holes by *Ae. albopictus* in the LNP probably implies complex interactions with native species, with possible species segregation. For example, in the LNP, we observed that *Ae. albopictus* relative abundance increased progressively with degree of anthropization, whereas conversely that of *Ae*. gr. Africanus decreased. Previous field and experimental observations in areas invaded by *Ae. albopictus* outside Africa clearly highlighted its aptitude to avoid interspecific competition and to coexist (or to displace) with resident North-American mosquito species laying eggs in tree holes, such as Ochlerotatus triseriatus (Juliano and Lounibos, 2005) and Ochlerotatus sierrensis (Kesavaraju et al., 2014; Washburn and Hartmann, 1992). Alternatively, *Ae. albopictus* might exploit water-filled rock-holes for its larval development (Pereira-dos-Santos et al., 2020). These microhabitats are extremely abundant in riverbeds or along the banks of streams in gallery forests in the northern part of the LNP. They also host a diversity of mosquito species communities (Philip, 1962), implying competitive interspecific interactions. The predation at natural larval development sites is an important issue that *Ae. albopictus* has to deal with because several Toxorhynchites species (e.g. *T. brevipalpis, T. viridibasis*) use tree holes and other phythotelmata in the LNP (C. Paupy, unpublished data, April 2019). Other potential resident predators (e.g. arthropods and nematodes) and locally circulating pathogens (i.e. entomopathogenic viruses, fungi, and parasites) are additional biotic constraints that *Ae. albopictus* might overcome to successfully colonize natural breeding sites in forests that are not free spaces. Abiotic factors that might slow down *Ae. albopictus* spread in the sylvatic niche are linked to the environment and to rainfall that modulates the filling and also the flooding of natural larval development sites. In the northern part of the LNP, the rainfall pattern includes two drought periods (4 and 2 months) leading to temperature increase, relative humidity decrease and the progressive drying of many natural water containers. During these periods, several mosquito species tend to concentrate in a few large natural water collections where interspecific competition is probably exacerbated, or survive as diapausing eggs in dried containers waiting for the next rains. *Aedes* albopictus possesses this trait because it is an ancestral character of *Stegomyia* species; however, resident sylvatic species with which it competes are probably tolerate better egg desiccation (Sota and Mogi, 1992). It has already been suggested that differences in thermal and desiccation tolerance of eggs are key environmental drivers of the spatial and temporal coexistence pattern between invasive *Ae. albopictus* and resident *Ae. aegypti* in Florida (Juliano et al., 2002). However, an experimental study demonstrated that desiccation resistance in non-diapausing eggs of *Ae. albopictus* is quickly selected (Sota, 1993). Moreover, during extreme rain spells, most rock-holes in riverbeds are rapidly flooded, causing important population crashes in mosquitoes that use this microhabitat. Overall, further field and experimental studies on the ecology of pre-imaginal *Ae. albopictus* in forested environments are necessary to improve the knowledge in larval ecology, to define its ecological limits, and to assess its population dynamics.

### Penetration pattern of *Aedes albopictus* inside forest ecosystems

To better understand the potential for ecological niche expansion into the primary forest of *Ae. albopictus* sylvatic populations in the LNP, it is crucial to determine its capacity to survive and reproduce beyond the forest margins that have been proposed as its preferred habitat (Fontenille and Powell, 2020). As our knowledge on the nature and abundance of natural larval microhabitats used by *Ae. albopictus* in African forests was limited, we used ovitraps to investigate its penetration and occupancy pattern of the forest ecosystem. The ovitrap survey carried out at three sites indicated a significant positive correlation between anthropization levels at the forest margins and *Ae. albopictus* relative abundance in ovitraps. By assuming that the availability of natural container breeding habitats was similar among sites, we postulate that the attractiveness of ovitraps was homogenous, and thus this result suggests that the *Ae. albopictus* population size (here approximated by egg density) remains strongly human-dependent even in a sylvatic ecological context. Although it is not strictly required for its occurrence, the human presence contribute to maintain *Ae. albopictus* density at the forest margins in the LNP, resulting in oviposition “hotspots” at these direct interfaces with humans (i.e. entry points), and confirming the close relation between *Ae. albopictus* and human settings from which females seems to move away only slightly. Anthropogenic water containers outside the forest could be the more productive larval development sites compared with natural ones (their biotic and abiotic constraints for *Ae. albopictus* have been discussed above). Moreover, even if *Ae. albopictus* displays some trophic plasticity and feeds off a large vertebrate host spectrum, humans remain among the most frequent hosts (Delatte et al., 2010; Pereira dos Santos et al., 2020). Besides humans, domestic animals, when available, are more frequent blood sources for *Ae. albopictus* collected deep in forests at anthropic-sylvatic ecotones in Brazil, compared to wildlife (Pereira dos Santos et al., 2018).

In addition to the effect on egg density, we observed a continuous decrease of laid egg density from forest margins (i.e. entry points) towards the deep interior of galleries. Indeed, most eggs were laid in the first 50 meters and then their number gradually decreased as the distance from the forest margin increased, becoming zero after 225 meters. In Brazil, where a similar study was carried out, the same oviposition pattern was observed, although *Ae. albopictus* eggs were detected deeper in the forest (up to 500 meters) (Pereira dos Santos et al., 2018). This difference could be related to the higher mosquito population size at the forest margin in Brazil, due to a greater density of human habitations (up to 10 habitations/km2 in Brazil versus 2 habitations/km2 at most in the inhabited compartment of the LNP). In their study, Pereira et al. observed many traces of human passage over several hundred meter deeper and deeper into the forest. Because of the natural tendency for *Ae. albopictus* to follow moving object in their quest of blood (Edman and Spielman, 2020), recurrent human movements in the forest might help *Ae. albopictus* to establish deeper in the forest. The occupancy pattern we highlighted, with several spatially consecutive positive spots, suggests that *Ae. albopictus* females gradually disperse from the margin into the forest ecosystem by successive short distances. Several laboratory and field experiments indicate that *Ae. albopictus* females practice skip oviposition that enhances its dispersal when looking for multiple oviposition sites during a single gonotrophic cycle (Davis et al., 2016, 2015; Fonseca et al., 2015). The maximum distance we observed (225 meters) from the forest fringe conforms with the reported distance range actively travelled by *Ae. albopictus* during a single gonotrophic cycle in human settings (Vavassori et al., 2019). Alternatively, dispersal deeper into the forest ecosystem could be related to other resource seeking behaviors, including blood and sugar meals and resting sites (Edman et al., 1998; Forattini, 1973). Overall, our results indicate that *Ae. albopictus* has a limited aptitude to colonize forest ecosystems beyond their margins, and suggest that a permanent establishment in forests, in the absence of human occurrence, would remain associated with low population densities, probably due to lower fitness in forest ecosystems, as previously discuss above. Regardless of population densities, *Ae. albopictus* presence at the forest margins raises the question of its interactions with the wildlife present in the LNP, and of its potential role in the transmission of enzootic arboviruses.

### *Aedes albopictus* sylvatic populations and the potential risk of enzootic arbovirus transmission to humans

*Aedes* albopictus propensity to evolve in natural ecosystems by exploiting natural larval breeding sites and feeding on different animal species increases the risk of animal-to-animal and animal-to-human transmission of zoonotic pathogens (Benedict et al., 2007; Delisle et al., 2015; Gubler, 2003; Paupy et al., 2012, 2009). Therefore, its colonization of natural ecosystems must be carefully monitored, especially in forested tropical areas where wildlife biodiversity and the risk of emerging infectious disease are high (Allen et al., 2017). *Ae. albopictus* spread in Central Africa, with the consequent replacement of the native species *Ae. aegypti*, has coincided with dramatic changes in the epidemiology of *Aedes*-borne viruses, with major outbreaks in cities and villages (Ngoagouni et al., 2015). In some cases, new viral variants particularly well adapted to be transmitted by *Ae. albopictus* have been locally selected and exported to other *Ae. albopictus* colonized areas with an increasing risk of epidemics (de Lamballerie et al., 2008). For example, in 2014, a CHIKV outbreak occurred in Montpellier (France) following its introduction from Cameroon (Delisle et al., 2015). Central Africa has become a source of *Ae. albopictus*-adapted arbovirus epidemic strains that can spread throughout and beyond Africa due to the increased flow of international travelers. The presence of *Ae. albopictus* in forest environments and its interactions with wild animals could further increase the risks of spillover and the emergence of *Aedes*-borne viruses from wildlife into the human compartment. Indeed, the occurrence of *Ae. albopictus* sylvatic populations in the LNP, sometimes in the absence of humans, implies that the species uses wildlife hosts as a source of blood, and might thus interact closely with the great diversity of enzootic viruses of the Congo Basin forests. Even if its spread were to be limited to forest fringes and forest galleries, interactions with wildlife would not still be limited. Indeed such environments gathering savanna and forest animals from diverse animal groups, are known having an increased animal diversity/density (Mares and Ernest, 1995; Redford and da Fonseca, 1986; Seaman and Schulze, 2010), offering thus to *Ae. albopictus* numerous opportunities to feed on and to pick-up enzootic viruses. This continuous block of tropical forest (the largest after the Amazonian Basin) is a hot-spot for enzootic cycles, including arboviruses that involve as vector sylvatic *Aedes* mosquitoes phylogenetically related to *Ae. albopictus* (i.e. all belonging to the *Stegomyia* subgenus) (Table 2). The LNP hosts a very abundant and highly diversified wildlife (>400 birds, 14 primates, 12 carnivores, 12 ungulates species, and >30 rodent and bat species) (Tutin, 1996; Vande Weghe, 2011) among which there are typical wildlife reservoirs of an extraordinary diversity of mosquito-borne enzootic viruses. For instance, nine species of monkeys present in the LNP (including 3 *Cercopithecus, 1 Cercocebus*, and *1 Colobus*) are natural reservoirs of yellow fever, CHIKV, ZIKV and other arboviruses. Contacts between *Ae. albopictus* and some of the sylvatic vertebrate hosts (at least at sites where humans or domestic animals are absent) opens up the opportunity for ongoing interactions with some of the enzootic arboviruses suspected to be circulating in the LNP (Table 1). Moreover, our findings that *Ae. albopictus* continues to bite humans entering these forested areas suggests a potential for a bridging role in the the horizontal transfer of enzootic viruses between wildlife and humans. This risk of transfer already exists in La Lopé region, as indicated by the presence of bridge vector species, such as *Ae. africanus*; however, none of them thrives as *Ae. albopictus* does in the human compartment. This means that a virus picked up from wildlife and transmitted (i.e. selected) to humans by *Ae. albopictus* at the forest interface would have an increased likelihood to produce secondary cases once introduced in human settings because already adapted to *Ae. albopictus*. This potential risk should be defined more precisely by complementary surveys, including field-collected mosquitoes, to identify the viruses carried by *Ae. albopictus* in sylvatic areas, and/or vector competence studies, to assess its ability to transmit sylvatic strains of viruses that circulate among its wildlife hosts in La Lopé region.

In conclusion, this study highlighted the penetration of the Asian tiger mosquito *Ae. albopictus* in the forest environment of Central Africa. We are witnessing a phase of invasion of the African forest ecosystem where sylvatic *Ae. albopictus* populations are mainly derived from human-associated populations. It remains to verify whether possible selection of more zoophilic lineages that can survive and reproduce in the absence of humans will eventually or has already occurred. From the One Health perspective, this introduction might increase the risk of transmission of zoonotic infectious agents for which *Ae. albopictus* could play the role of bridge vector, as previously hypothesized by some (Fontenille and Powell, 2020; Gubler, 2003; Pereira dos Santos et al., 2018). It is now crucial to determine *Ae. albopictus* capacity to exploit natural breeding sites in forest ecosystems (e.g. tree and rock holes) and to feed on wild vertebrates, and the effect of this invasion on the life cycles of sylvatic pathogens.

## Methods

### Study area

The LNP (11.61303E, 0.10409S) covers 4,964 km^2^ (Henschel et al., 2005) of equatorial forest and is among the most important biodiversity hotspots in Gabon, with very abundant and diversified wildlife (Vande Weghe, 2011). The northern part of the LNP is dominated by a forest-savanna mosaic landscape crossed by the Ogooué River and several smaller waterways along which La Lopé village is located (Figure 1). In this village with <2,000 inhabitants (UNESCO, 2006), the economic activity is essentially based on wildlife conservation and ecotourism, with a large number of visitors from Gabon and other countries (∼1,000 visitors per year), coming by road or by train (DGT-Gabon, 2018). Further south, the forest-savanna mosaic landscape gives way to dense equatorial forest with several forest-savanna ecotones.

### Study design and sampling procedure

The different sampling periods corresponded to the long rains because the LNP has a bimodal rainy season (long rainy and short rainy periods, from February to May and from October to November, respectively) interspersed with dry periods. First, to explore the range of *Ae. albopictus* dispersal from La Lopé village towards the wild compartment of the LNP, we carried-out a sampling session using ovitraps and BG-Sentinel traps in May 2014. For ovitrap-based sampling, 35 spatial spots were randomly selected in the village (n=10) and in the wild compartment southwards (n=25) (Figure 1). Collection sites were at least 300 m apart and each contained five ovitraps left active for 14 consecutive days. Traps consisted of sections of bamboo trees (15-20 cm long and 5-7 cm in diameter) that contained 300 mL of tap water and a 15 × 5 cm wood paddle for egg laying. We hung these traps on tree trunks at 1m-1.5m from the ground. In the wild compartment of the park, we placed traps along the network of forest galleries that connects the village to the forest block, and also in some selected isolated forest patches and at forest fringes (Figure 1). We deployed individual BG-Sentinel traps equipped with BG-lure and a carbon dioxide source (mixture containing 22g of yeast, 2kg of saccharose and tap water to a final volume of 5L) from 8 am to 6 pm in the village (i.e. anthropic compartment; 1,340 Trap x hours of sampling effort) and in the wild side of the LNP (i.e. sylvatic compartment; 1,645 Trap x hours of sampling effort). To follow the dynamics of *Ae. albopictus* invasion inside the park, we completed the dataset with additional occurrences reported for the species from 2017 and 2018 during *Aedes* aegypti surveys (Aubry et al., 2020; Xia et al., 2020, 2018). These observations were obtained using several sampling methods, including larval prospections in natural water collection sites (e.g. rock pools, tree holes), domestic and peri-domestic water containers, bamboo or plastic ovitraps, BG-Sentinel traps, and human landing catches.

Second, to assess *Ae. albopictus* colonization level in the forest ecosystem (i.e. its “capacity to exploit forest environments for suitable oviposition sites”) (Pereira dos Santos et al., 2018), in April 2019, we carried out a field experiment in the village and wild compartment of the park. Three sampling sites were selected to assess whether the human footprint (or human presence) degree influences the colonization of forests ecosystems by *Ae. albopictus*. The sites were selected and spatially distributed based on a relative anthropization gradient: “inhabited forest edge” (*WCS*: in the village, a section of 30 neighboring habitations), “sparsely inhabited forest edge” (*Station 1*: forest base camp, less than 5 human habitations housing 2-10 people who permanently live or work at this site site), and “non inhabited forest edge” (*Station 2*: no human habitation) (Figure 1). In this study, “anthropization” refers to human presence or frequentation, evidenced by human dwellings or traces of human passage or activities. We considered that a site was anthropized (or inhabited) when there was at least one human habitation within ≤300 m from the collection site. At the interfaces between anthropic and forest compartments (*WCS* and *Station 1*) and between savanna and forest (*Station 2*), we deployed ovitraps from the fringe towards the deeper parts of the forest compartment. We surveyed the colonization by *Ae. albopictus* by monitoring along the transects the number of eggs laid by females in ovitraps placed at different distances from the edge towards the forest. At each site, we set ovitraps at the intersections of a grid of parallel lines spaced 25m apart and crossed perpendicularly by parallel lines spaced 25m apart. At each site, the grid extended from the edge (0 m) to 150 m (*WCS*), 125 m (*Station 1*), and 175 m (*Station 2*) inside the forest compartment, depending on the spatial conformation of the forest blocks (Figure S2). Ovitraps consisted of dark plastic cups that contained 300 mL of mango leaf infusion and a 15 cm x 5 cm hardboard paddle for egg collection. In total, we deployed 105 ovitraps (35 at *WCS*, 30 at *Station 1*, and 40 at *Station 2*). At each site, we monitored oviposition for 10 consecutive days with a change of paddles after 5 days. We transported the recovered paddles to the insectary for 3 days of drying and egg counting. Then, we immersed wood paddles separately in 1 L distilled water in a plastic tray (40 × 30 × 7 cm) for egg hatching. After 3 days of immersion, we removed the paddles from the trays and air-dried them at room temperature for 3 additional days before immersion in new distilled water following the same procedure. We fed mosquito larvae 0.01 g *TetraMin* ® fish food (Spectrum Brands, Inc.) every day and reared them to the adult stage. We morphologically identified adult mosquitoes using “custom” taxonomic keys based on updates of the Edwards’ identification keys for Ethiopian mosquitoes (Edwards, 1941) and the Huang’s key for the subgenus *Stegomyia* of *Aedes* mosquitoes from Afrotropical Region (Huang, 2004), and then counted them for statistical analysis.

### Statistical analysis

All statistical analysis were carried out using the R software, v3.6.1 (https://www.r-project.org/). Assessment of *Ae. albopictus* dispersion from the village towards the wild compartment of the park was based on its presence at collection sites. To assess the level of colonization of the forest ecosystem, we used count data from the transect experiment. Because the number of emerged adult mosquitoes was not equal to the number of collected eggs in the field due to egg viability, we used the weighted number of *Ae. albopictus* eggs (WNAE) as the main variable of interest. The WNAE is an estimation of the number of *Ae. albopictus* eggs collected relative to the percentage of *Ae. albopictus* adults after emergence. We calculated the WNAE as the frequency of *Ae. albopictus* adults multiplied by the total number of eggs collected in the field for each collection site. Then, we tested various generalized linear models to fit the penetration into the forest by *Ae. albopictus* using the “WNAE” as the response variable, and the “Distance” of the trap to the forest edge and the “Habitat” as fixed and random effects, respectively. We did not include data from *Station 2* in the models because of insufficient number of positive traps. We checked the WNAE distribution normality visually and using the Shapiro-Wilk normality test. We tested different models with a log link function and two alternative error structures: Poisson or negative binomial, using the stats, glmmADMB, lme4, and MASS packages (Bates et al., 2014; Fournier et al., 2012; Ripley et al., 2013). Then, we used the predict function for model visualization and interpretation. We used the Akima package to interpolate in two-dimensions the spatial distribution of the density of *Ae. albopictus* eggs in ovitraps (Akima et al., 2016).

## Acknowledgments

We thank all study participants. We are grateful to La Lopé National Park and all staff members, especially Mr. Josué Edzang-Mve for his precious assistance in the field experiment implementation. Special gratitude to the staff of the laboratory *Écologie des Systèmes Vectoriels* (ESV) of the Interdisciplinary Center for Medical Research of Franceville (CIRMF, Gabon): Dr Boris Makanga, Dr Martha Nigg, Dr Neil Michel Longo-Pendy, Mr Patrick Yangari and Mr Ousmane Akone-Ella.

## Research Authorization

This study was authorized by the National Center of Scientific and Technical Research (CENAREST, Gabon) under the authorization N° AR0010/19/MESRS/CENAREST/CG/CST/CSAR and AR0020/14/CENAREST/CG/CST/CSAR.

## Authors’ contributions

Paper writing: JON, CP

Experiment conception and design: JON, DA and CP

Field investigation: JON, MFN, CP

Material identification: JON, CP

Technical advisors: NR, DJ, PK, CC

Data analysis: JON, DR

General supervision: CP

## Figure legends

**Figure S1:**
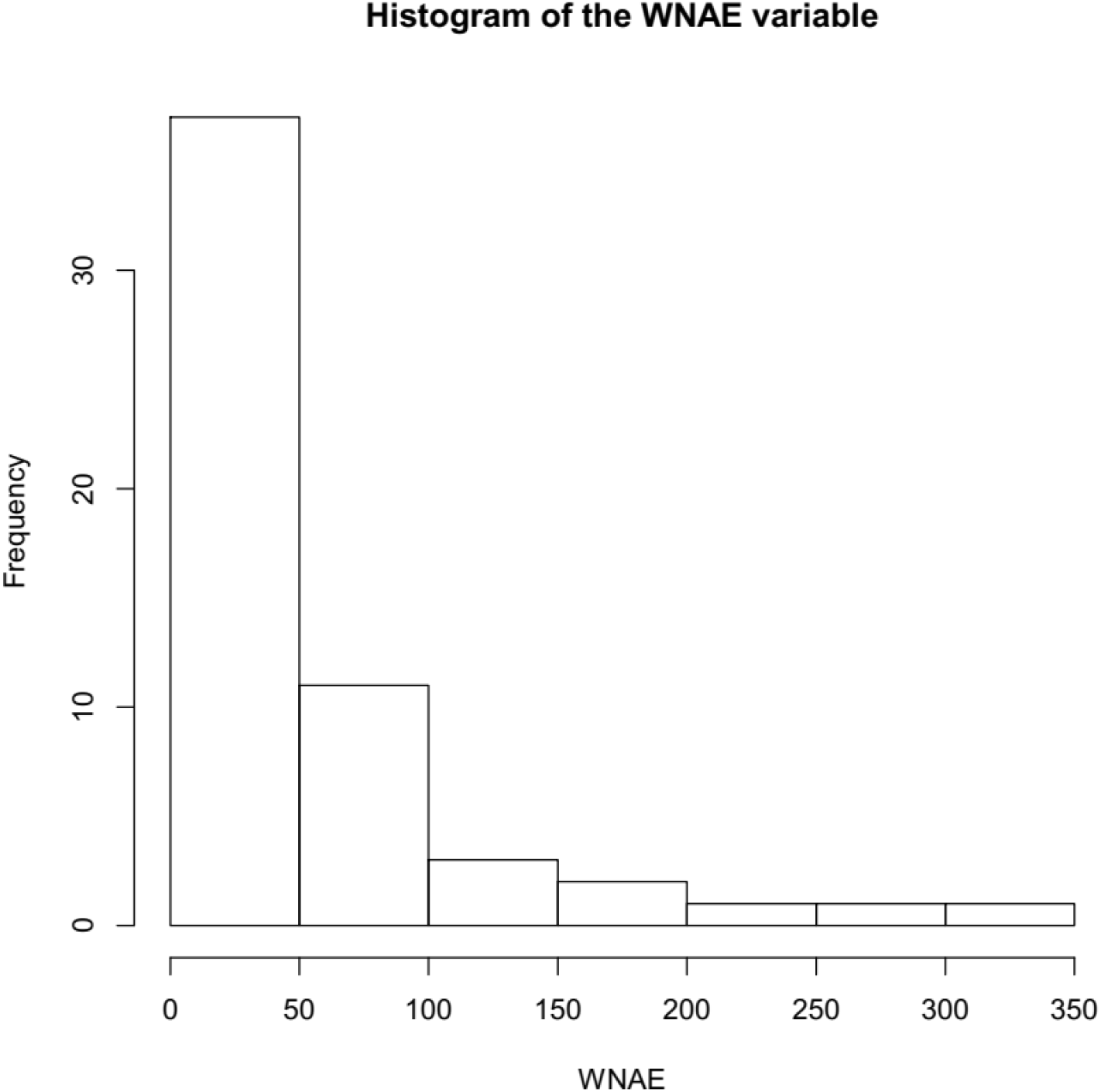
Histogram showing the distribution of the weighted number of *Ae. albopictus* eggs (WNAE), highlighting its non-normal distribution.

**Figure S2:**
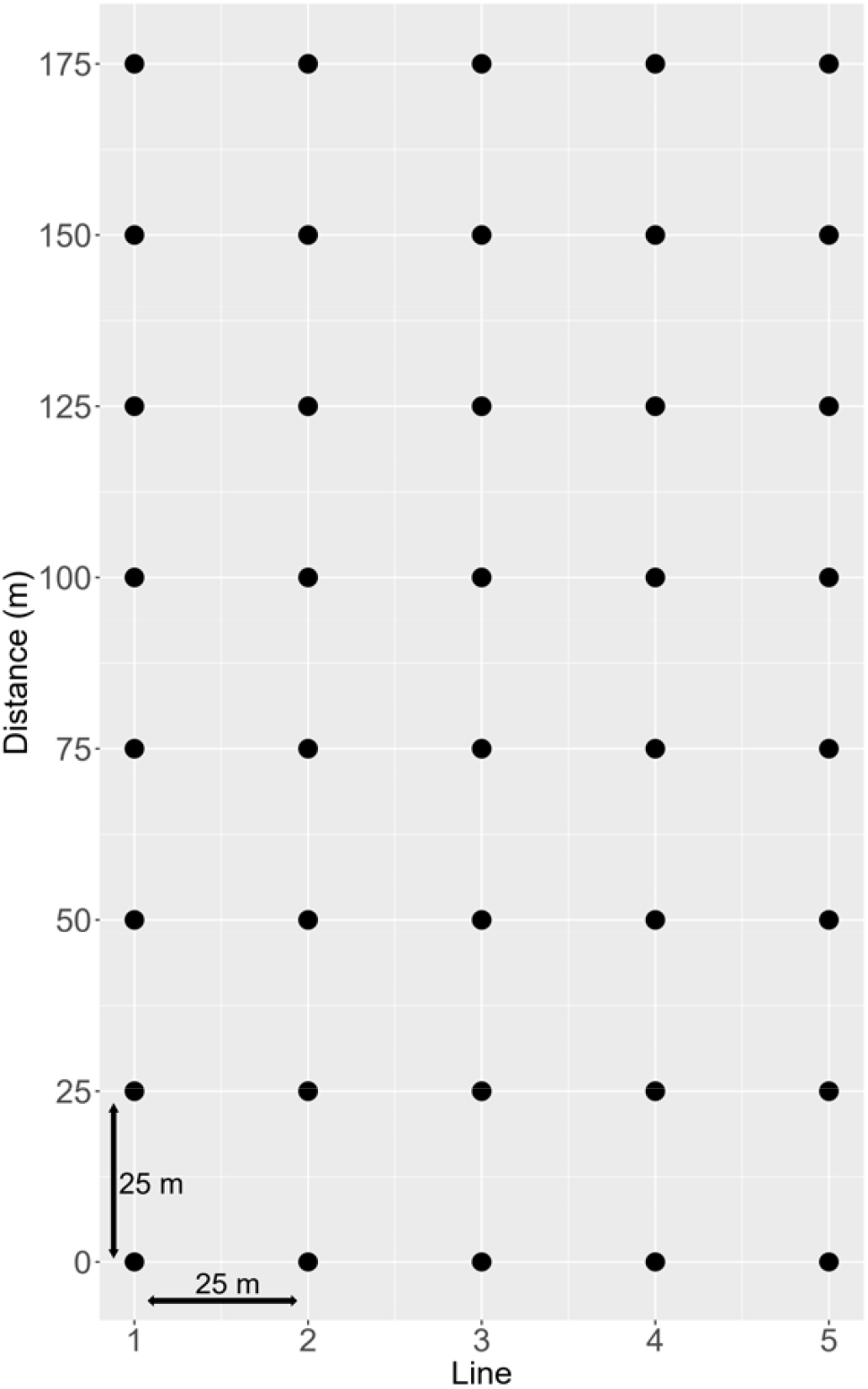
Sampling plan diagram for ovitrap-based experiments implemented along anthropization gradients at *WCS, Station 1*, and *Station 2* to assess *Ae. albopictus* penetration pattern inside forest ecosystems. Black full circles represent the position of individual ovitraps placed following five lines from the forest edge to the forest interior. The maximum distance from the forest edge varied according the site and the spatial conformation of the forest block (150 m for *WCS*, 125 m for *Station 1*, and 175 m for *Station 2*).

